# Single-Cell Transcriptomes and Immune Repertoires Reveal the Cell State and Molecular Changes in Pemphigus Vulgaris

**DOI:** 10.1101/2023.04.13.536499

**Authors:** Shumin Duan, Qionghua Li, Fei Wang, Wenjing Kuang, Yunmei Dong, Dan Liu, Jiongke Wang, Wei Li, Qianming Chen, Xin Zeng, Taiwen Li

## Abstract

The etiology and pathogenesis of pemphigus vulgaris (PV) are closely related to both immune cells and epithelial cells, but the specific subtypes of immune cells involved in PV and their roles are not yet fully understood. Additionally, the specific functions and mechanisms of first-line treatment glucocorticoids on cell types of PV remain to be elucidated. We performing 5’ single-cell RNA sequencing, combined with V(D)J enrichment on buccal mucosal lesions and peripheral blood samples from treatment-naïve patients with PV, in conjunction with post-treatment peripheral blood samples obtained after oral prednisone treatment. Our findings suggest that IL-1α signaling pathway, myeloid antigen presenting cells, inflammatory CD8+ Trm, and dysfunctional CD4+ Treg are crucial in PV. Our results were also supported by immunohistochemical assays. Furthermore, our results show that prednisone has a significant impact on monocytes and MAIT, but a limited effect on CD4+ Treg. Finally, we provide CDR3 amino acid sequence data of BCR that may be used as therapeutic targets. In conclusion, this study provides a comprehensive understanding of PV, particularly in the mucosal-dominant type, and the effect of GCs on PV, which could effectively lead to the development of new therapeutic strategies.

## INTRODUCTION

Pemphigus is a rare, chronic autoimmune bullous disease that poses a significant threat to human health. Pemphigus vulgaris (PV), which comprises 60%-90% of all cases[1–3], is characterized by autoantibodies that cause the separation of keratinocytes, resulting in flaccid blisters and hemorrhagic erosions of the skin and mucous membrane[3]. The binding of autoantibodies to desmoglein (DSG) 3 is believed to play a crucial role in the development of the disease. Research has shown that targeting common sequences in the complementarity-determining region 3 (CDR3) of pathogenic autoantibodies could be an effective way of preventing or treating the disease[4]. Additionally, autoantibodies alter the activity of various signaling pathways, including Ca^2+^, p38MAPK, PKC, Src, EGFR/Erk etc., that regulate cell cohesion and desmosomal component turnover[5], suggesting that targeting these pathways may also be beneficial for patients[4].

Autoimmune diseases such as PV are thought to arise from a combination of genetic and environmental factors that activate antigen-presenting cells (APCs). These APCs subsequently educate autoreactive T cells by presenting specific Dsg peptides via their HLA class II molecules[3]. Several subtypes of immune cells have been found playing various roles in the pathogenesis of PV. For example, helper T cells (Th), CD8+ T cells, and γδ T cells are mainly pathogenic[6–8]. Conversely, regulatory T (Treg) cells mediate immunosuppression or tolerance, and the decrease in Treg number or their dysfunction can promote the pathogenesis of PV[9–15]. As immunology continues to develop, more and more types of immune cells with different functions have been discovered, and it is essential to clarify their roles in PV. Furthermore, given that the oral mucosa is the primary target site of PV’s autoantibodies, studying the interactions between epithelial cells, immune cells, and their role in oral mucosal lesions is crucial for gaining a deeper understanding of PV’s pathogenesis.

Glucocorticoids (GCs) are widely used in clinical settings as an anti-inflammatory treatment and are the first-line treatment for PV[3]. Prednisone is often the preferred GC for treating PV. In 1948, Philip Hench and Edward Kendall discovered the anti-inflammatory and immunomodulatory effects of GCs[16]. Follow-up studies found that GCs primarily achieve their anti-inflammatory effects by inhibiting the activity of T cells, NK cells and other immune cells[17–19]. The immunosuppressive and anti-inflammatory effects of GCs are mainly mediated by the GC receptor (GR), which directly represses proinflammatory transcription factors (TFs)[20, 21]. However, the actions of GCs appear to be cell-type and disease context-dependent[22–24], and the specific cell types and intracellular targets of GCs in PV are not yet identified. To address this gap, we apply single-cell RNA sequencing (scRNA-seq) combined with T cell receptor (TCR) and B cell receptor (BCR) sequencing to characterize the cellular changes for PV. We present a comprehensive single-cell immunological landscape and identify immune cell types and their expression patterns of functional genes that might contribute to PV development. Furthermore, we examined the effects of prednisone treatment on the peripheral immune system of PV patients, focusing on the regulation of cell type-specific gene expression and TF activity. Our results support the established roles of various T cell types, B cells, myeloid cells, and epithelial cells in the pathophysiology of PV and the effects of prednisone at the single-cell level. Additionally, our findings reveal the clonal features of T and B lymphocytes in PV.

## MATERIAL and METHODS

### Experimental design

The objective of this study was to characterize the cellular composition of PV’s buccal mucosal lesions and peripheral blood at a single-cell resolution. The study design was performing 5′ single-cell RNA sequencing, with V(D)J enrichment for select samples.

### Patient recruitment and sample collection

The subjects of this study were from the patients who were diagnosed as mucosal-dominant type of PV in the Department of Oral Medicine [25], West China Hospital of Stomatology, and the healthy controls were from those who planned to undergo orthognathic surgery in the Department of Orthognathic Surgery, West China Hospital of Stomatology. Selection criteria of PV included (1) age ranging from 18 to 65 years; (2) diagnosed as PV, and the diagnosis criteria was based on recommendations of an international panel of experts [26]. The diagnoses were confirmed with clinical manifestations, histology, Dsg-specific antibody tests and immunohistology criteria. Selection criteria of healthy controls included (1) age ranging from 18 to 65 years; (2) oral mucosa is heathy and oral mucosal diseases are excluded after consultation and examination. The sample collection was approved by the Institutional Review Board of West China Hospital of Stomatology. All the participants gave their consent by signing informed consent. Buccal mucosa biopsy samples from PV patients were obtained pre-treatment. Buccal mucosa samples of healthy controls were collected during orthognathic surgery. Peripheral blood from PV patients were obtained pre-treatment and when lesions cured after a period of oral prednisone treatment. During surgical resection, specimens were placed in normal saline and immediately maintasined on ice pending further processing. Clinical characteristics are summarized in Additional file 1: Table S1.

### Single-cell suspension preparation

Oral mucosa samples were processed with the least delay after collection. Each tissue sample was minced and then digested using the human Whole Skin Dissociation Kit (Miltenyi Biotec; No. 130-101-540) at 37`℃`. The digested sample was filtered through a 70 μm cell strainer (Miltenyi Biotec; No. 130-110-916) to remove large particles. The filtered mixture was centrifuged and the supernatant was removed. The remaining cell deposition was resuspended in Red Blood Cell Lysis Buffer (Abcam; ab204733) for 5 min to lysing erythrocytes. After washing and resuspending in D-PBS(BBI Life Science; E607009-0500), the cell suspension was processed using magnetic cell separation (MicroBeads, Buffers, Columns, Separators: Miltenyi Biotec) to sort living cells.

Fresh blood samples were collected in EDTA tubes, diluted with D-PBS, and then transferred onto Ficoll-Paque isolation solution (GE Healthcare;17-5442-02). PBMCs were slowly and carefully collected after density gradient centrifugation, and then added with Red Blood Cell Lysis Buffer for 5 min. Afterward, PBMCs were washed twice with D-PBS.

### Library construction and sequencing

Library construction of scRNA and VDJ was conducted using Chromium Single Cell V(D)J Reagent Kits (v1 Chemistry,10X Genomics) following the manufacturer’s instructions. All sample libraries were sequenced on a Nova-Seq6000 sequencer (Illumina, USA).

### Single-cell RNA-seq data analysis

The FASTQ files from each sample were processed through Cell Ranger v7.0.0 pipeline from 10X Genomics using GRCh38 genome reference. The output files include gene expression and TCR/BCR data. Then, we used the R package Seurat v4.2.0 to analyze aforementioned single-cell gene expression data. All individual Seurat objects from the same tissue (PBMCs or buccal mucosal tissue) were integrated using the merge function. In the quality control phase, we filtered data based on the percentage for transcripts of mitochondrial genes, the number of unique molecular identifiers (UMI) and the number of genes detected. Each filtered Seurat object was then normalized using Seurat’s NormalizeData function. Highly variable genes were detected using the FindVariableFeatures function in Seurat. The “ScaleData” function is used to linearly transform the data, that is, to scale the expression of each gene so that the difference between cells is 1. Principle component analysis (PCA) was performed. We use “Harmony” package to remove the batch effects when necessary. Clusters were then identified using Uniform Manifold Approximation and Projection (UMAP).

### Sub-clustering of T/NK, myeloid cells, and epithelial cells

T/NK, myeloid cells, and epithelial cells were extracted from buccal mucosal tissues for further sub-clustering. After extraction, genes were scaled to unit variance. PCA and clustering were performed as described in Dimension Reduction and Major Cell Type Annotation section. A very small number of contaminated cells in the extracted subsets are discarded.

### CCell function scoring

The function AddModuleScore in the Seurat package was used to compute the scores of functional modules. We chose some gene sets in the GO term of Homo sapiens as functional modules, including T cell activation involved in immune response (GO:0002286), B cell activation involved in immune response (GO:0002312), and Myeloid cell activation involved in immune response (GO:0002275).

### Differential gene expression and functional enrichment analysis

Differential gene expression analysis was conducted by applying the FindMarkers function in Seurat with default parameter settings. The R package clusterProfiler was applied to functional enrichment analysis. Gene lists arose from biological process (BP) terms in the GO Resource.

### Cell-cell interaction analysis

To comprehensively analyze cell-cell interactions between immune cells, we used CellChat (v 1.5.0). We derived potential ligand-receptor interactions based on the expression of a receptor by one cell subpopulation and ligand expression by another.

### Transcription Factor-Target Gene Network Analysis

Based on the gene regulation identified in our scRNA-seq data, we utilized the GENIE3 R packages (v 1.18.0), as well as the RcisTarget database (v 1.16.0) of the SCENIC (v 1.3.1) workflow to predict the transcription factor and their downstream genes. We used GENIE3 to computerize the genetic regulatory networks from our expression data. We further used RcisTarget databases to recognize the enriched transcription factor-binding motifs and those potential downstream genes (regulons).

### TCR/BCR clonotype analysis

TCR clonotypes were identified as VDJ genes of TCR α-β paired chains combined with the nucleotide sequence of CDR3 regions. Similarly, BCR clonotypes were defined as the VDJ genes of paired light and heavy chains combined with the nucleotide sequence of the CDR3 region. The R package scRepertoire was utilized in TCR and BCR analysis. The Upset plot was implemented by the UpSetR package. The Chord diagram was made by circlize package.

### Immunohistochemistry (IHC) and Multiplex immunofluorescence

Paraffin 4-μm-thick sections were prepared. DAB Detection Kit (Polymer) (GK600705, Gene Tech) was used for IHC staining. Multiplex immunofluorescence staining was performed by using the Opal Polaris five-color immunohistochemistry staining kit (Akoya Biosciences). The following primary antibodies were used: anti-human IL-1 alpha (ab300501, Abcam), anti-human CD8 (MA5-14548, Thermofisher), and anti-human CD69 (ab233396, Abcam).

### Statistical analysis

All of the statistical analyses were conducted using software R v4.2.0. The detailed descriptions of statistical methods are stated in the section of results.

## RESULTS

### Activated immune characteristics in PV buccal mucosal lesions

To explore the cell composition and molecular characteristics of oral mucosal lesions and peripheral circulation in PV, we carried out scRNA-seq combined with TCR/BCR sequencing on samples obtained from buccal mucosal lesions and peripheral blood of two patients with PV. We also successfully obtained peripheral blood samples from both PV patients when their lesions were cured after oral prednisone treatment and performed the same sequencing protocol. Paired samples were collected from healthy donors, consisting of two oral buccal mucosa tissues and four peripheral blood samples (Fig. 1a). The clinical pictures and histopathological examination results are shown in Fig. 1b. Additional details regarding the clinical characteristics of the donors are provided in Table S1 and Fig. S1a.

**Fig. 1.**
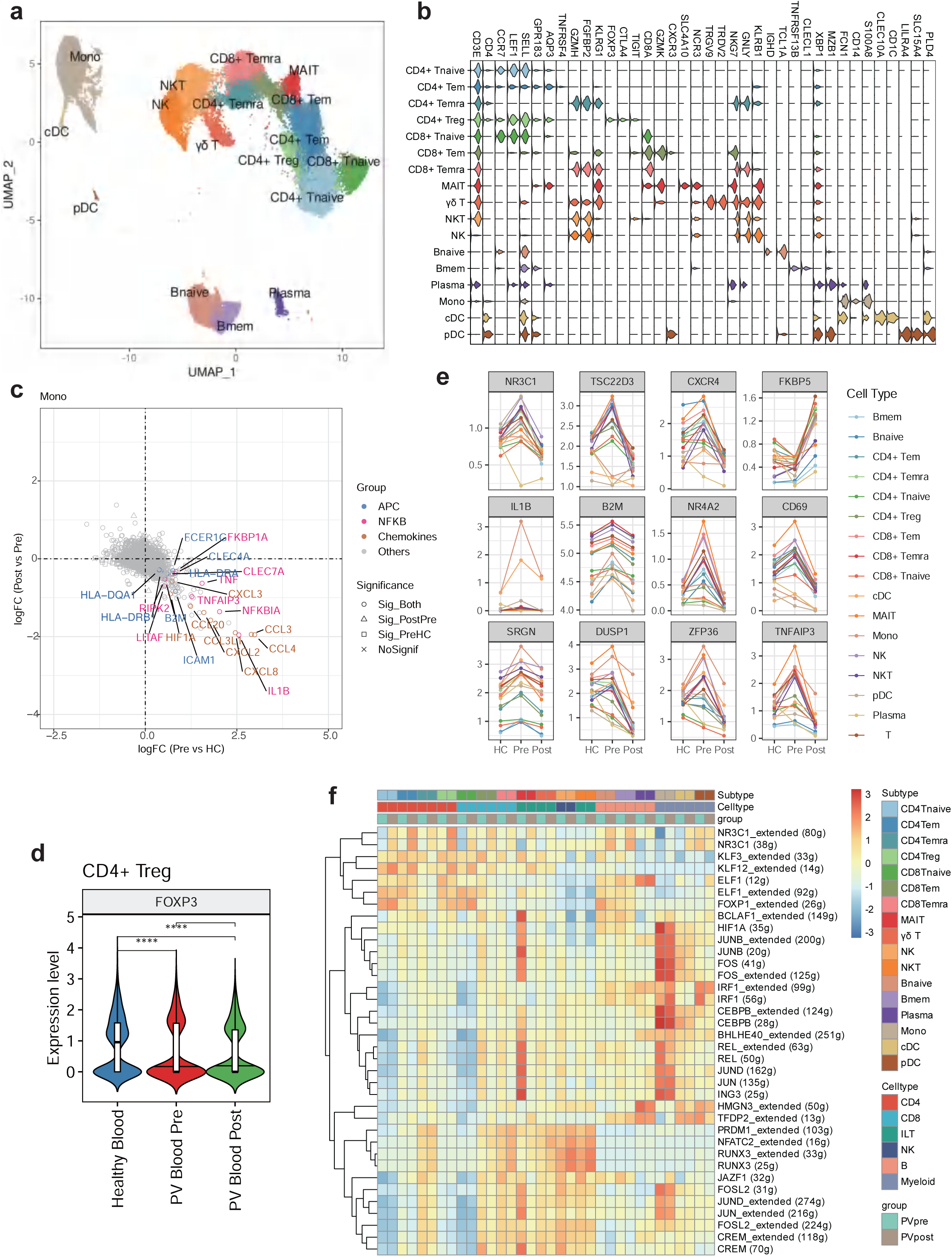
Activated immune characteristics in PV buccal mucosal lesions. **a** The experimental design of scRNA and scVDJ sequencing (PV Tissue(n=2), PV Blood Pre(n=2), PV Blood Post(n=2), Healthy Tissue(n=2), Healthy Blood(n=4)). **b** Light microscopy images of 2 patients’ biopsy specimen from the buccal mucosa showing supra-basal blister formation. Direct immunofluorescence microscopy of a perilesional biopsy specimen shows intercellular deposits of IgG and C3 in the epidermis. Clinical pictures show 2 patients had erosions affecting buccal mucosa and the lesions were cured after oral prednisone treatment. **c** All of the cells were grouped into 11 types. The bottom figures presents cells’ distribution in HC and PV group. **d** Heatmap shows classical gene markers of each cell population. **e** Cell type proportion in healthy tissues and PV tissues. **f** Barplots exhibit the related scores of different three immune cell subtypes.

After quality control, we obtained 35,309 cells for all tissue samples, with 13,764 cells were from healthy controls (HCs) and 21,545 cells from PV lesions. We performed unsupervised clustering and uniform manifold approximation and projection (UMAP) analysis on these cells (Fig. 1c). Cells were initially classified into 11 main cell populations using canonical markers, including 4 immune cell types: T/NK cells (CD3E,XCL1,XCL2), B cells (MS4A1, CD79A, CD83), plasma cells (JCHAIN, IGKV3-20, IGKV1-5), myeloid cells (HLA-DRA, HLA-DPA1, IFI30), and 7 non-immune cell types: fibroblasts (CFD, DCN, PTGDS), myofibroblasts (TAGLN, ACTA2, MYL9), epithelial cells (KRT14, S100A2, KRT15), endothelial cells (TM4SF1, ACKR1, CCL14), melanocytes (DCT, PMEL, TYRP1), myocytes (ACTA1, CKM, MYLPF), mast cells (TPSAB1, TPSB2, CPA3) (Fig. 1d). We observed more T/NK cells, B/Plasma cells and myeloid cells in PV lesions compared with HCs, which is consistent with previous findings[27–31] (Fig. 1e). T/NK cells were the main immune cells in the both groups of tissue samples, followed by myeloid cells, while B cells and Plasma cells accounted for the least.

Furthermore, to understand functional states of above immune cell types, we selected relevant genes from the Gene Ontology (GO) database and scored their expression levels in three immune cell types. Compared with HCs, T/NK cells and myeloid cells in PV lesions exhibited a more activated immune status, while B cells were more prone to differentiate to plasma cells (Fig. 1f). Collectively, our data showed overactivated immune characteristics in PV buccal mucosal lesions.

### Multiple signaling pathways have increased in adjacent epithelium of PV erosions

As PV is an organ-specific autoimmune disease that primarily affects epithelial cells[1, 27], we first examined the epithelial cell subsets and identified a total of 2246 epithelial cells after performing quality control. After batch correction (Fig. S1b), we identified four subtypes on the basis of their marker gene expression. (Fig. 2a). Basal cells were characterized with expression of COL17A1 and KRT19[32, 33], as COL17A1 encodes the alpha chain of type XVII collagen which is a structural component of hemidesmosomes at the dermal-epidermal basement membrane (Fig. 2b). Proliferating cells were identified as they expressed markers of cell proliferation (MKI67, TOP2A, STMN1)[34]. Superfacial cells expressed genes involved in differentiation (CLDN4, SPRR1B, SBSN)[35, 36]. As expressing higher KRT13 than basal cells and without superfacial gene markers[37], suprabasal cells were identified. What’s more, we found DSG3 was expressed in cells in each layer, while DSG1 was mainly expressed in superfacial layer cells (Fig. 2c). This finding was consistent with current knowledge and supported our identification of cell types[25].

**Fig. 2.**
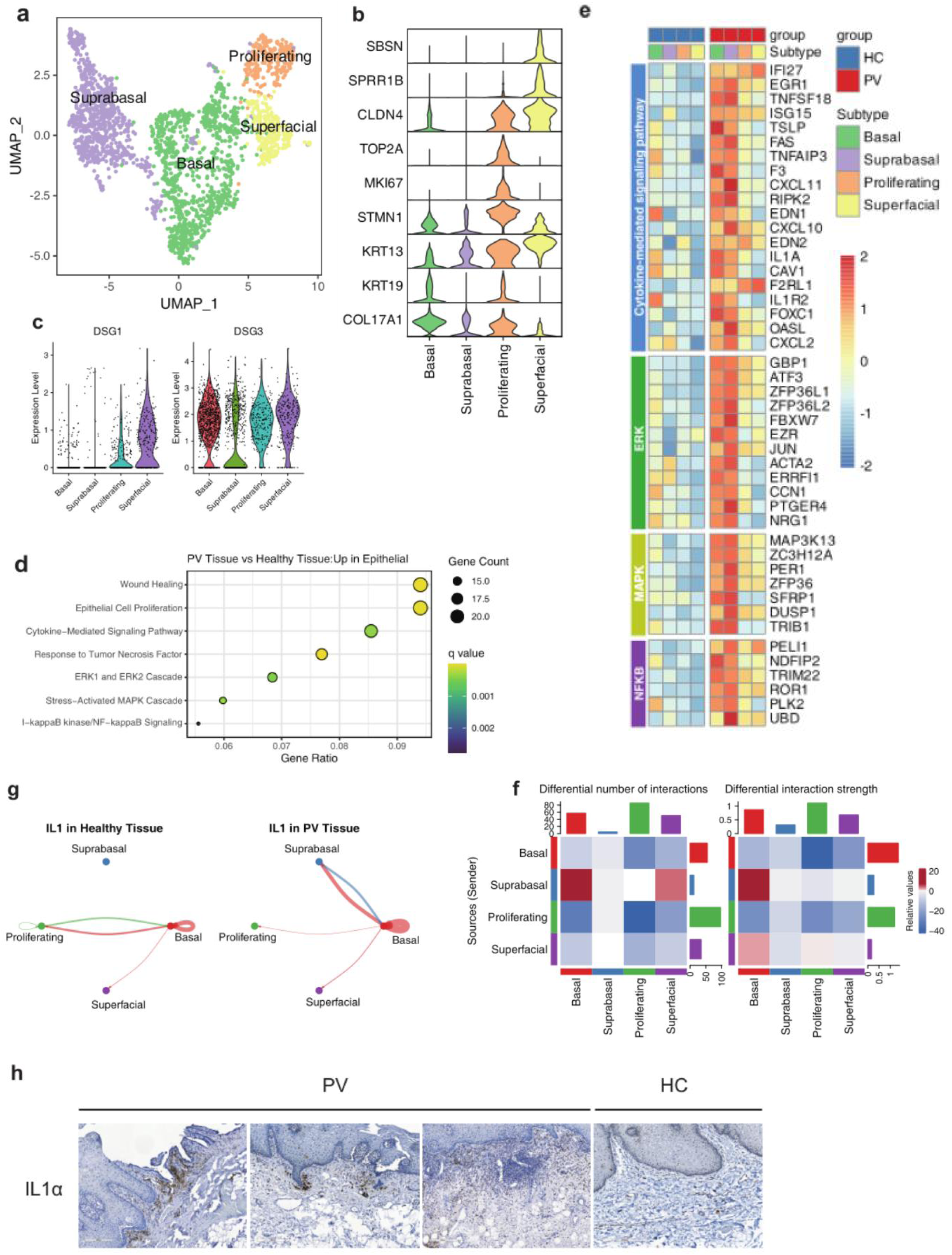
Multiple signaling pathways have increased in adjacent epithelium of PV erosions. **a** Epithelial cells were grouped into 4 types. **b** Violin plot shows classical gene markers of 4 epithelial cell subtypes. **c** Violin plot shows expression of DSG1 and DSG3 in 4 epithelial cell subtypes. **d** Dot plot reveals GO enrichment terms of genes with differential expression (average log_2_^FC^(PVs vs HCs) > 0.5 and adjusted p value < 0.05). **e** Heatmap shows gene expression pattern of 4 epithelial cell subtypes between the two groups. **f** Heatmaps shows differential number and strength of interactions among 4 epithelial cell subtypes. Positive value (color red) represent increasing number and strength of interactions in PVs compared with HCs. Negative value (color blue) represent decreasing number and strength of interactions in PVs compared with HCs. **g** Circle plots show the inferred network of the IL1 signaling pathway in the two groups. Edge width indicates the inferred communication probabilities. **h** Immunohistochemistry of IL-1α expression in the buccal mucosal lesions of 3 PV patients and 1 healthy donor.

Additionally, we found that adjacent epithelial cells of PV erosions upregulated pathways related to “wound healing” and “epithelial cell proliferation”, indicating a repairing and proliferating response of the adjacent epithelium to PV erosions (Fig. 2d). Moreover, several signaling pathway such as “cytokine-mediated signaling pathway”, “response to tumor necrosis factor”, “ERK1 and ERK2 cascade”, “stress-activated MAPK cascade” and “ I-kappaB kinase/NF-kappaB signaling” have increased in adjacent epithelium of PV erosions, which was consistent with previous studies [38–42]. The heatmap showed the related genes involved in the signaling pathways (Fig. 2e). We observed a general upregulation of related genes in epithelial cells of PV group, especially in basal and suprabasal layers. Since histopathology of PV reveals suprabasal split formation, the above signaling pathways and related genes are highly associated with acantholysis in PV. Moreover, in the “cytokine-mediated signaling pathway”, we could see FAS, interleukin-1a (IL1A), TNFSF18, and TNFAIP3 were upregulated in perilesional PV mucosa, which were consistent with previous reports[43–45]. By comparing the number and strength of cell interactions among epithelial cells, we found an enhanced interaction between basal layer and suprabasal layer (Fig. 2f), and the IL1 signaling was significantly upregulated between the two layers (Fig. 2g). We performed immunohistochemical staining to explore the expression patterns of IL-1 alpha in the buccal mucosal lesions of PV. We observed increased infiltration of IL-1 alpha in the buccal mucosal lesions of PV patients (Fig. 2h). Therefore, local inhibition of IL-1 or the above signaling pathways may hold potential as a new targeted therapy for PV.

### Gene expression pattern of myeloid subclusters

As myeloid cells play an important role in the early stage of PV’s pathogenesis through antigen processing and presentation, we extracted myeloid cells (excluding mast cells) for further analysis. After removing a few polluted cells, we obtained 2642 cells and identified 5 subtypes (Fig. 3a, Fig. S1c). Based on the presence of monocyte markers (FCN1, AQP9, IL1RN, LILRA5) and the absence of macrophage markers (MRC1, PLTP, MSR1), monocytes grouped apart from macrophages[46] (Fig. 3b). Macrophages could be further divided into CCL2(high) and MAF+ macrophages. CCL2, also known as monocyte chemoattractant protein-1 (MCP-1), is well known for its ability to drive the chemotaxis of myeloid and lymphoid cells[47], while MAF is important for macrophage terminal differentiation [48].

**Fig. 3.**
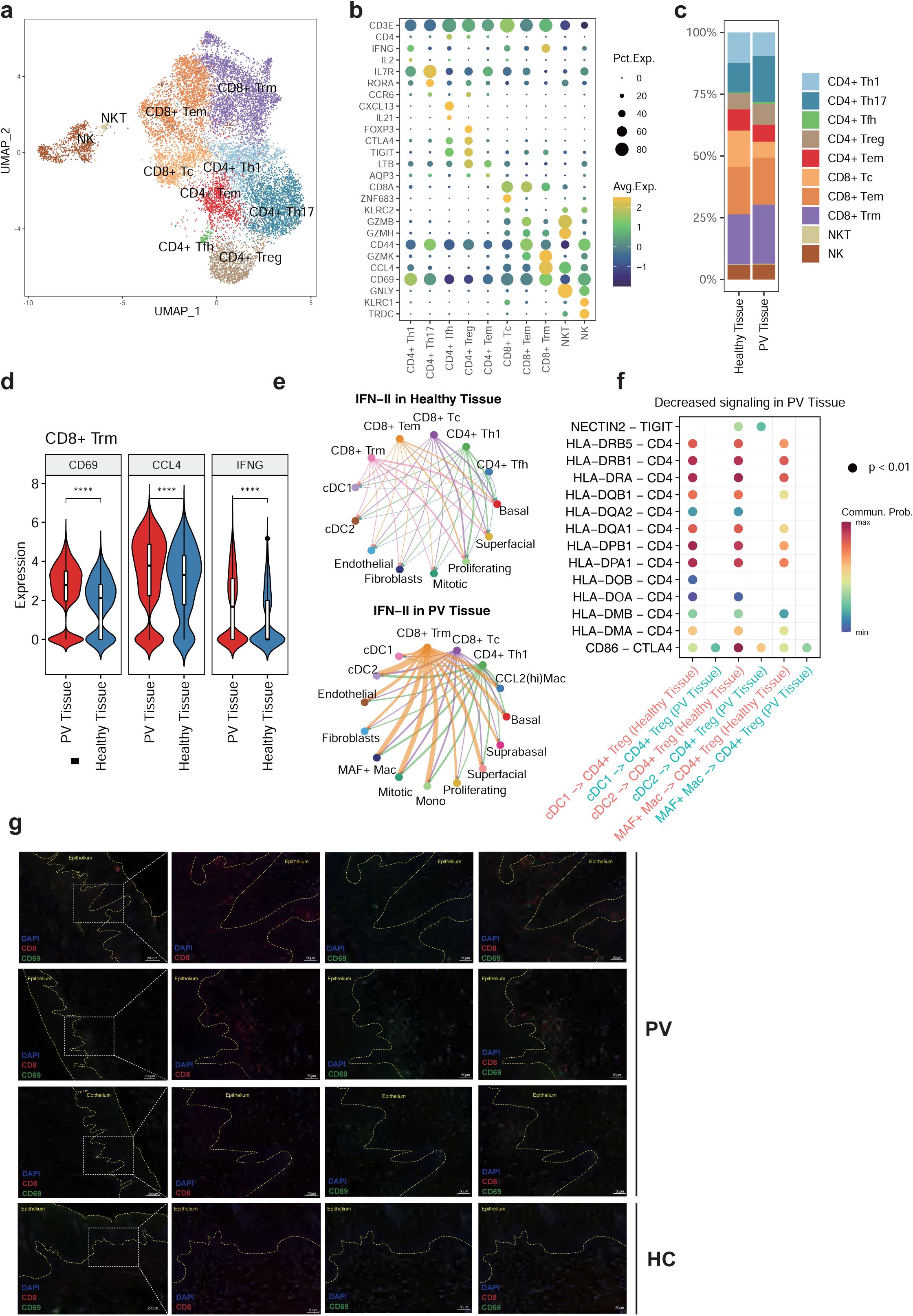
Gene expression pattern of myeloid subclusters. **a** Myeloid cells were grouped into 6 types. **b** Violin plot shows classical gene markers of 6 myeloid cell subtypes. **c** Heatmap shows gene expression pattern of 6 myeloid cell subtypes. **d** Pseudotime trajectory analysis of Mono/Mac(Mono, CCL2(hi)Mac, MAF+ Mac), cell density plots are displayed along x and y axes. Color-coded for the Mono/Mac phenotypes (left panel), the group (middle panel) and the pseudotime (right panel). **e** Density plots reflect the number of cells along the 3 Mono/Mac subtypes stratified for HCs and PVs. **f** Profiling of chemokine genes along the trajectory to confirm CCL2(hi)Mac’s functional annotation. **g** Gene expression dynamics along the Mono - CCL2(hi)Mac - MAF+Mac linage. Genes group into 3 distinct gene sets, each of which has distinct expression profiles.

Dendritic cells (DCs) were also divided into cDC1 (CLEC9A, XCR1, NET1) and cDC2 (CLEC10A, CD1C, CD1E, FCGR2B). Moreover, a small group of cells expressing proliferating markers (MKI67, PCNA, UBE2C) was identified as mitotic cells. As the heatmap shows (Fig. 3c), different cell types have different patterns of gene expression. Monocytes highly express genes related to chemotaxis. CCL2(hi)Mac also expressed chemokine genes but without expressing monocytes markers (S100A8, S100A9)[46]. MAF+ Mac highly expressed genes in the complement system and antigen-presentation relevant (major histocompatibility complex) MHC-I/II genes. cDC1 cells were characterized by high expression of classical MHC-I genes (HLA-A/B/C) and MHC-II genes, whereas cDC2 cells highly expressed MHC-II genes. The gene expression patterns of cDC1 and cDC2 were consistent with their different roles in driving particular T cell responses[49]. cDC1 is mostly distinguished by the cross-presentation of intracellular antigens to CD8+ T cells and the direct presentation of antigens to CD4+ T cells. cDC2 is effective at directly presenting extracellular antigens to CD4+ T cells and eliciting humoral immune responses of type 2 and type 17. As the gene expression signature of CCL2(hi)Mac is between Mono and MAF+ Mac, we utilized trajectory analysis to clarify the potential differentiation among them (Fig. 3d-g). We identified CCL2(hi)Mac cluster as an intermediate cell population that acts as a chemoattractant for monocytes and other immune cells by expressing various chemokines [50] (Fig. 3f, g). MAF+ Mac cells in PV were enriched towards the end of the cell lineage (Fig. 3e). These results highlighted the heterogeneity of myeloid cells in the buccal mucosa and suggested functional specialization in specific subpopulations.

### Myeloid subclusters have activated antigen processing and presentation

DCs are professional antigen-presenting cells that play a critical role in regulating the balance between immunity and tolerance, and changes in the specialized DC system are a common feature of both systemic and tissue-specific autoimmune diseases [51, 52]. We observed a significant increase in the proportion of both cDC1 and cDC2 in PV lesions compared to HCs (Fig. S1d). The higher number of cDCs in PV lesions could potentially contribute to the loss of tolerance observed in PV.

As monocytes are no longer viewed only as precursor cells for macrophages, and their emerging roles as antigen-presenting cells have been reviewed [53], we found that Mono and MAF+ Mac upregulated MHC-I/II genes (Fig. 4a,c). We further performed GO analysis with these genes (Fig. 4b, d), and found that both cell types upregulated pathways related to T and B cell activation, as well as antigen processing and presentation.

**Fig. 4.**
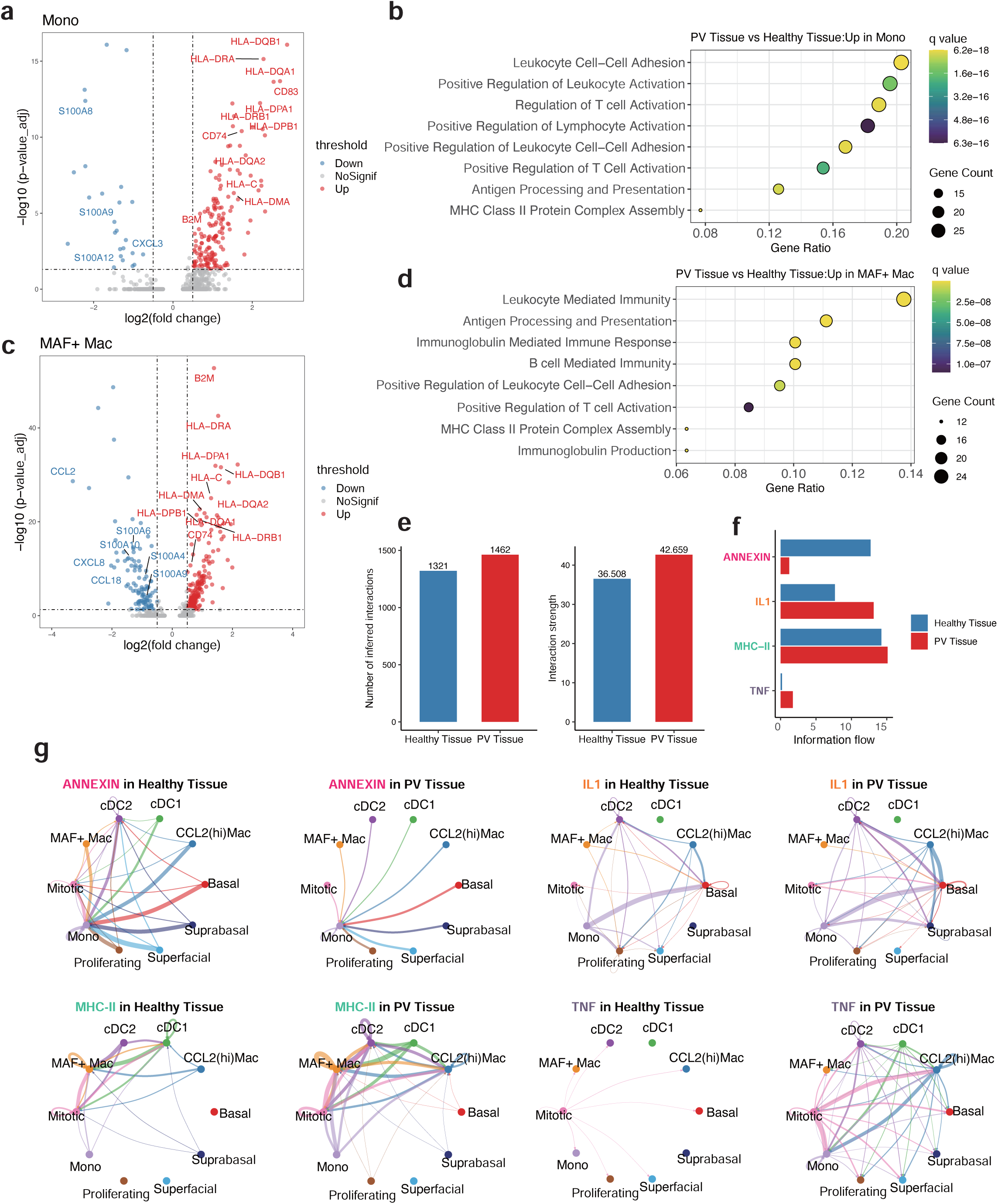
Myeloid subclusters have activated antigen processing and presentation. **a, b** Volcano plot shows the differentially expressed genes of monocytes(Mono)(**a**) and dot plot reveals GO enrichment terms of genes with differential expression (average log_2_^FC^(PVs vs HCs) > 0.5 and adjusted p value < 0.05)(**b**). Wilcoxon Rank Sum test was used. **c, d** Volcano plot shows the differentially expressed genes of MAF+Mac(**c**) and dot plot reveals GO enrichment terms of genes with differential expression (average log_2_^FC^(PVs vs HCs) > 0.5 and adjusted p value < 0.05)(**d**) in PV tissues in comparison with HCs. Wilcoxon Rank Sum test was used. **e** Bar plots reflects the number and strength of cell interaction along epithelial cells and myeloid cells in the two group. **f** The bar graph displays the selected signaling pathways in the comprehensive information flow of PV tissues and healthy controls. **g** Circle plots show the inferred network of the ANNEXIN, IL1, MHC-II and TNF signaling pathways in the two groups. Edge width indicates the inferred communication probabilities.

Moreover, we observed an increase in both the number and strength of interactions between myeloid and epithelial subclusters in PV lesions (Fig. 4e). We further analyzed specific signaling pathway networks and found that “Annexin” was decreased in PV (Fig. 4f, g), which could potentially impede its anti-inflammatory effects locally [54]. As we observed IL-1 alpha infiltration in subepithelial tissues in PV lesions (Fig. 2h), we also saw “IL1” signaling pathway among myeloid cells and epithelial cells was increased (Fig. 4h). Our results were consistent with the point of view from a recent review [55]. They pointed out tissue damage would trigger release of intracellular IL-1α from dying cells, thus generating an IL-1α-containing milieu, which is sensed by infiltrating myeloid cells expressing the IL-1R1. And they also considered IL-1α a central role in the pathogenesis of numerous conditions characterized by organ or tissue inflammation. Based on our findings, IL-1α is also involved in the pathogenesis of PV. Moreover, “MHC-II” and “TNF” were increased in PV (Fig. 4g), suggesting that blocking these signaling pathways could be potential treatment strategies. In summary, our findings suggest that myeloid subclusters play a key role in initiating and promoting PV by activating antigen processing and presentation in PV lesions.

### Inflammatory CD8+ Trm and selectively incompetent Treg in PV lesions

We next focused on T/NK cell subsets, which account for the largest proportion of immune cells in the tissue. By extracting T/NK cell population (14026 cells) from the whole cell map and correcting batch (Fig. S2a), we identified multiple functional T/NK cell subsets using a combination of previously reported markers and differentially expressed genes (DEGs), namely CD4+ helper type 1 T cells (CD4+ Th1; IFNG, IL2), CD4+ helper type 17 T cells (CD4+ Th17; IL7R, RORA, CCR6), CD4+ follicular helper T cells (CD4+ Tfh; CXCL13, IL21), CD4+ regulatory T cells (CD4+ Treg; FOXP3, CTLA4, TIGIT), CD4+ effect memory T cells (CD4+ Tem; LTB, AQP3), CD8+ cytotoxic T cells (CD8+ Tc; ZNF683, KLRC2, CD44^low^), CD8+ effect memory T cells (CD8+ Tem; GZMB, GZMH, CD44^hi^), CD8+ tissue-resident memory T cells (CD8+ Trm; GZMK, CCL4, CD69^hi^), natural killer T cells (NKT; GNLY, GZMB, GZMH), and natural killer cells (NK; CD3E^-^, GNLY, KLRC, TRDC) (Fig. 5a, b). We found an increasing proportion of CD4+ Th17 and CD4+ Tfh in PV lesions (Fig. 5c), which is consistent with a previous study [28]. There is also an increasing proportion of CD8+ Trm. And we also had immunohistochemical evidence (Fig. 5g). Besides, CD8+ Trm in PV lesions up-regulated the expression of CD69, IFNG and CCL4 compared with healthy controls (Fig. 5d). We further found the IFN-II signaling pathways between CD8+ Trm and other cell types were significantly upregulated in PV lesions (Fig. 5e). These findings suggested that CD8+ Trm cells were activated and played a significant part in the PV local immune response.

**Fig. 5.**
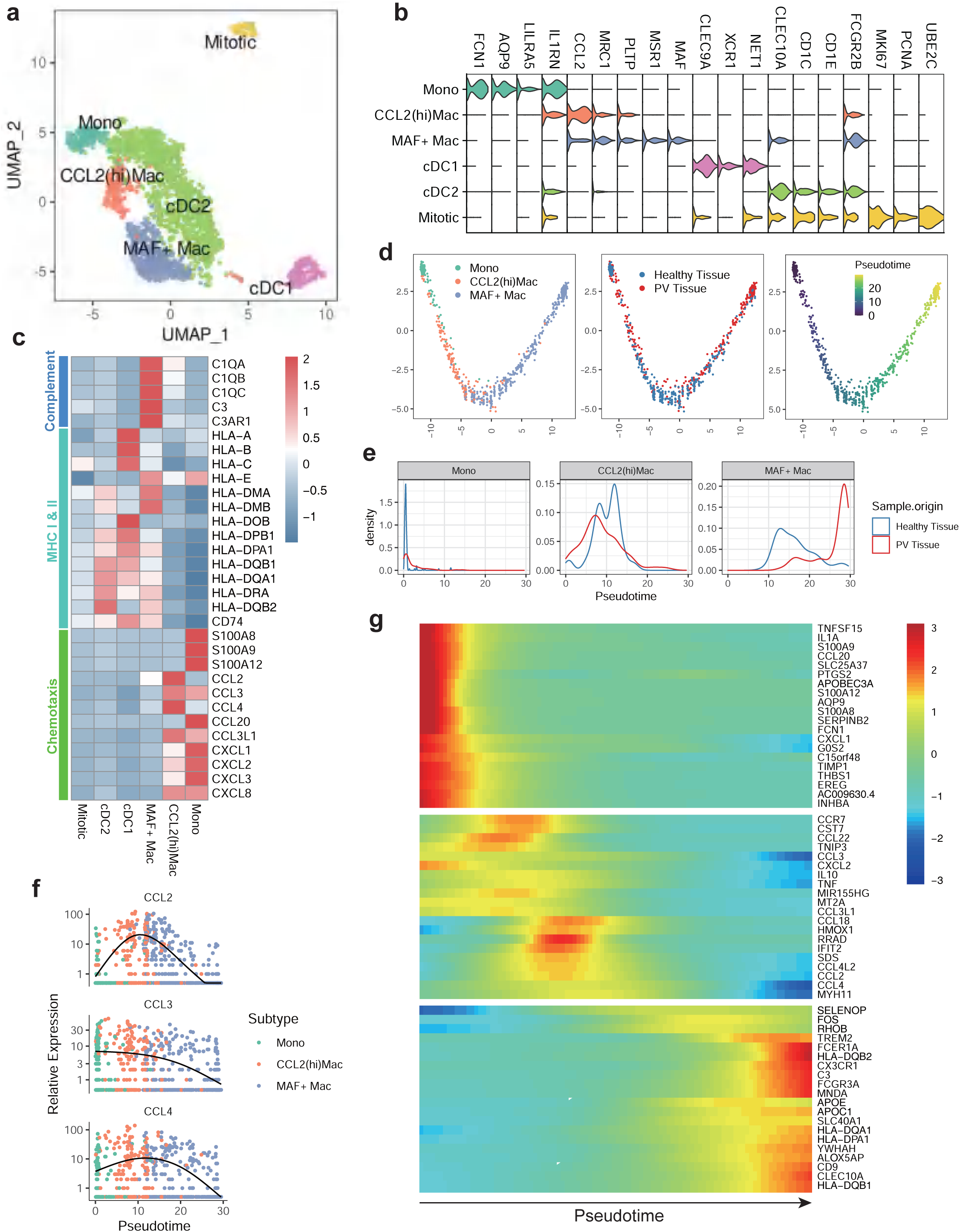
Inflammatory CD8+ Trm and selectively incompetent Treg in PV lesions. **a** T/NK cells were grouped into 10 types. **b** Dot plot shows classical gene markers of 10 T/NK cell subtypes. **c** Proportion of T/NK cell subtypes in healthy tissue and PV tissue. **d** Violin plots show the expression of CD69, CCL4 and IFNG in CD8+ Trm in the two groups. Wilcoxon Rank Sum test was used. **e** Circle plots show the inferred network of the IFN-II signaling pathways in the two groups. Edge width indicates the inferred communication probabilities. **f** Bubble plot shows selected decreased signaling ligand-receptor pairs between CD4+ Treg and myeloid subclusters in PV tissues compared to the HCs. **g** Multiplex immunofluorescence assay of CD8 (red), CD69 (green), and DAPI (blue).

The proportion of CD4+ Treg also increased in PV lesions. It is consistent with the results of the study of rheumatoid arthritis (RA), which showed Treg is enriched in RA synovial fluid[56–64]. However, by comparing the interaction between CD4+ Treg and other immune cells, we found the inhibitory signaling CD86-CTLA4 was down-regulated between Treg and myeloid subclusters (cDC1, cDC2 and MAF+ Mac), and NECTIN2-TIGIT was down-regulated between Treg and cDC2 (Fig. 5f). This reduction in Treg inhibitory signaling to myeloid cells may be a key factor contributing to the development of PV. It is possible that the enrichment of Tregs in PV lesions may represent a compensatory mechanism of the immune system, aimed at compensating for their lack of function and exerting their ability to inhibit the proliferation and cytokine secretion of certain immune cells, ultimately helping to control the local immune response.

### Prednisone affects cell type specific gene expression and transcriptional regulatory networks of PBMCs in PV

After data preprocessing and quality control, we obtained single-cell transcriptomes of 66,846 immune cells from PBMCs. Of these, 16,409 cells were from PV patients before treatment, 13,381 cells from PV patients when all lesions cured after oral prednisone treatment, and 37,056 cells from HCs. Using unsupervised graph-based clustering, we identified 17 clusters in PBMCs (Fig. 6a, Fig. S2b), which consisted of four types of CD4+ T cells: CD4+ naïve T cells (CD4+ Tnaive; CCR7, LEF1, SELL), CD4+ effector memory T cells (CD4+ Tem; GPR183, AQP3, TNFRSF4), CD4+ terminally differentiated effector memory T cells (CD4+ Temra; GZMH, FGFBP2, KLRG1), and CD4+ regulatory T cells (CD4+ Treg; FOXP3, CTLA4, TIGIT), three types of CD8+ T cells: CD8+ naïve T cells (CD8+ Tnaive; CCR7, LEF1, SELL), CD8+ effector memory T cells (CD8+ Tem; GPR183, GZMK, CXCR3) and CD8+ terminally differentiated effector memory T cells (CD8+ Temra; GZMH, FGFBP2, KLRG1), three types of innate-like T lymphocytes (ILT): mucosal-associated invariant T cells (MAIT; SLC4A10, NCR3), natural killer T cells (NKT; CD3E+, CD4-, CD8-, NKG7, GNLY, KLRB1) and γδ T (TRGV9, TRDV2), three types of B cells: naive B cells (Bnaive; IGHD, TCL1A), memory B cells (Bmem; GPR183, TNFRSF13B, CLECL1) and plasma cells (Plasma; XBP1, MZB1), three types of myeloid cells: monocytes (Mono; FCN1, CD14, S100A8), conventional/classical dendritic cells (cDC; CLEC10A, CD1C), and plasmacytoid dendritic cells (pDC; LILRA4, SLC15A4, PLD4) and natural killer cells (NK; CD3E-, NKG7, GNLY, KLRB1) (Fig. 6b).

**Fig. 6.**
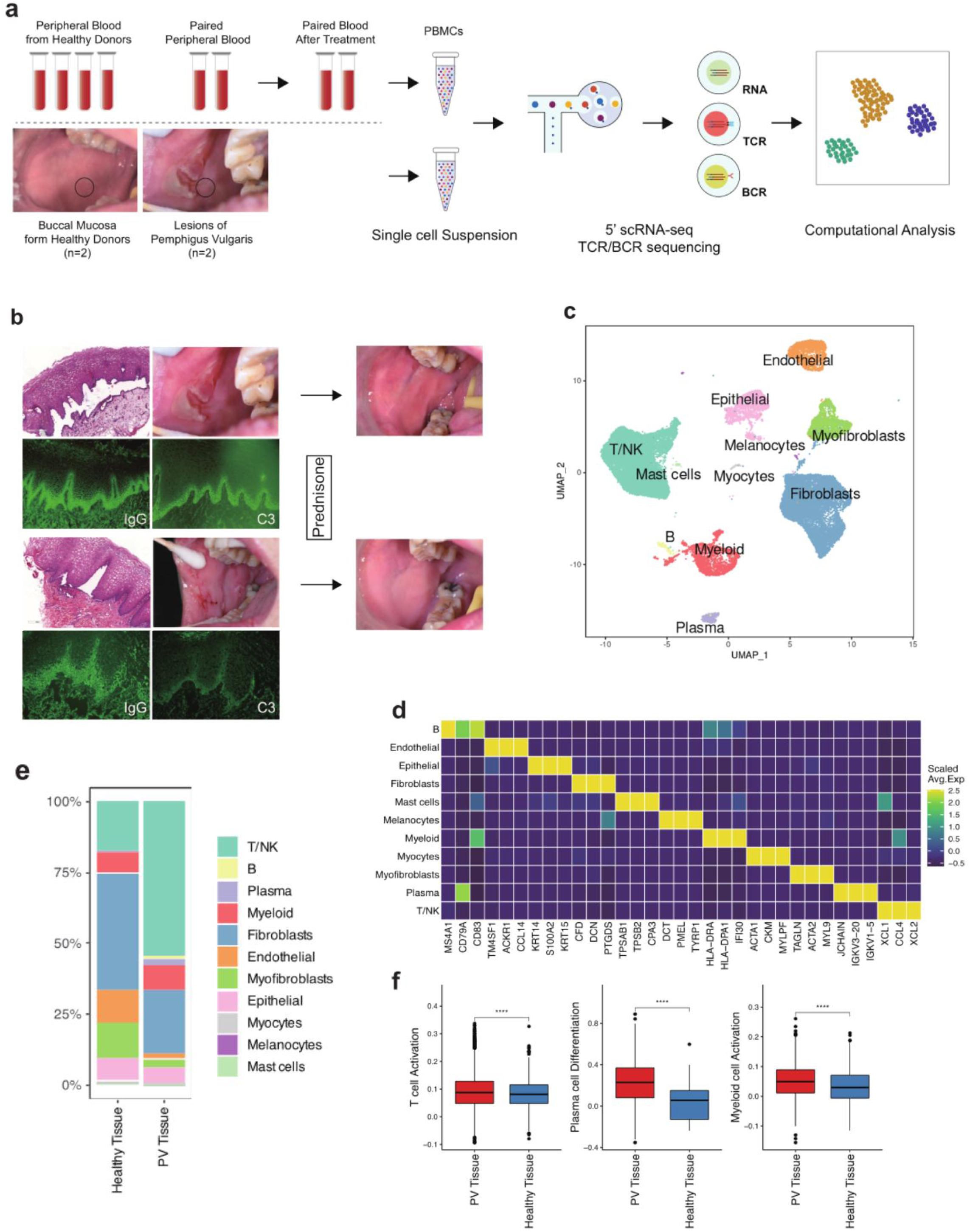
Prednisone affect cell type specific gene expression and transcriptional regulatory networks of PBMCs in PV. **a** Immune cells in PBMCs were grouped into 17 types. **b** Violin plot shows classical gene markers of 17 cell types in PBMCs. **c** Differential expression genes, comparing PV blood before treatment (Pre) vs healthy blood (HC) and PV blood after treatment (Post) vs PV blood before treatment (Pre), showing antigen presentation, NFκB and chemokines related genes were greatly up-regulated in PV blood before treatment (Pre) compared to HC, and down-regulated after treatment(Post). **d** Gene expression changes of critical genes among groups. **e** Gene FOXP3 expression alteration in CD4+ Treg in three groups. **f** Heatmap shows TFs down-regulated in Post vs Pre using down-regulated differential expression genes(average log_2_^FC^(PVs vs HCs) > 0 and adjusted p value < 0.05). APC: antigen presentation cells related, HC: healthy controls, Pre: PV blood before treatment, Post: PV blood after treatment.

To explore the cell type-specific gene expression characteristics of immune cells in peripheral blood of PV patients and to compare the changes after oral prednisone treatment, we found monocytes to be the cell type of noteworthy alterations. As Fig. 6c shows, the up-regulated genes in PV before treatment compared with HCs (Pre vs HC) were significantly down-regulated after oral prednisone treatment (Post vs Pre). The genes were chemokines, NFKB and antigen presentation-related genes. GO analysis revealed that the biological processes about monocytes activation were up-regulated in PV before treatment, and these biological processes were all successfully down-regulated when lesions cured after prednisone treatment (Fig. S2c). As CD4+ Treg’s critical role in autoimmune diseases and FOXP3 is required for Tregs’ development, maintenance and function[65, 66], we compared the expression levels of FOXP3 between the groups and found that it was lower in PV before treatment than in HCs and did not increase after prednisone treatment.

We also found some genes that were reversed after treatment in most cell types (Fig. 6e). NR3C1, known as glucocorticoid receptor (GR) gene, was down-regulated after treatment. It is suggested that the long-term exogenous GCs treatment would leave most mononuclear cells in the blood of PV patients in an anti-inflammatory state, as NR3C1’s expression would not be necessary when the cell is in an anti-inflammatory or suppressed state [67]. And it could possibly be one of mechanisms of GCs resistance. Moreover, TSC22D3, the gene encoding GILZ (glucocorticoid-induced leucine zipper) in almost all cell types was also down-regulated after treatment. As GR can inhibit NF-κB indirectly by inducing the expression of TSC22D3 [21], the gene’s down-regulation could be viewed as a result of NR3C1’s down-regulation. Besides, CXCR4 gene expression was up-regulated in PV’s blood and down-regulated after treatment, which was consistent with one previous scRNA study [68]. FKBP5 gene expression was up-regulated after treatment. It was also found in healthy bronchial biopsies following budesonide inhalation[69] and 10 patients with autoimmune Addison’s disease after intravenous infusion of 100 mg hydrocortisone[70]. IL1B was up-regulated in myeloid subclusters (Mono, cDC, pDC) and down-regulated after treatment. Combining our findings in PV lesions, IL1 related genes especially IL-1B and signaling were generally activated in local lesions and systemic circulation of PV. As a result, local or systemic use of IL-1 inhibitors has the potential to become one of the primary or adjunctive treatments for PV. There was a general increase of MHC-I related gene (B2M) expression in all cell type especially myeloid subclusters compared with HCs, and we saw a decrease after treatment.

The nuclear receptor 4A2 (NR4A2) was found up-regulated in the peripheral blood T cells of an autoimmune disease multiple sclerosis (MS) and viewed as an essential transcription factor for triggering the inflammatory cascade of MS[71, 72]. Our data also showed the same results, and the function and roles of NR4A2 in PV need further study. Other genes showing the same changes included T cell activation associated gene CD69[73], and inflammation related gene SRGN[74, 75]. DUSP1, ZFP36 and TNFAIP3 were down-regulated after treatment. The results were contrary to the current knowledge that GCs activate the above genes and related signaling pathways to play an anti-inflammatory role. The reasons for the variations could be due to different research subjects and methods, as well as the action of glucocorticoids being disease and cell-type-specific.

Given the anti-inflammatory action of GR mainly depends on its ability to directly repress proinflammatory transcription factors[21, 76], we applied SCENIC to predict the critical TFs that could interact with GR. Our results showed that down-regulating TFs included glucocorticoid receptor NR3C1, AP-1 components (JUN, JUNB, JUND, FOS and FOSL2), and Kruppel like factor family (KLF3 and KLF12) (Fig. 6f). Moreover, we found that AP-1 components (JUN, JUNB, JUND and FOS), REL, one member of nuclear factor κB (NF-κB) family, was highly active in both MAIT and monocytes before treatment, indicating MAIT and monocytes in peripheral blood of PV had an activated inflammatory status. In addition, monocytes highly expressed CEBPB which is a key transcription factor regulating monocytic gene expression. After treatment, the activity of those regulons in MAIT was significantly decreased, and MAIT had the sharpest change in the gene expression mentioned above, indicating that MAIT is an important target cell type for the anti-inflammatory effect of prednisone.

### The characteristics of TCR and BCR in PV

To investigate the clonal features of T cells among different sample groups, we analyzed TCR sequences in combination with scRNA-seq data. In PV01 patient’s lesion, we found that cells with clonotype overlap with those in peripheral blood were CD8+ Trm (Fig. 7a). This cluster of cells was in a state of hyperexpanded clonal status (Fig. 7b,c), while the cells in peripheral blood were CD8+ Tem (Fig. 7d, Fig. S3a). Similar findings were observed in PV02 (Fig. S3b-d). The clone intensity was higher in lesions than in peripheral blood (Fig. 7c, Fig. S3a-c). Additionally, we found that the TRAV38-2DV8/TRAJ34 pair of alpha chain and TRBV19/TRBJ2-1 pair of beta chain were preferentially used in TCRs from PV01 lesion (Fig. 7e), and the cells expressing TRAV38-2DV8 and TRBV19 were overlapped with the cells described above (Fig. 7f).

**Fig. 7.**
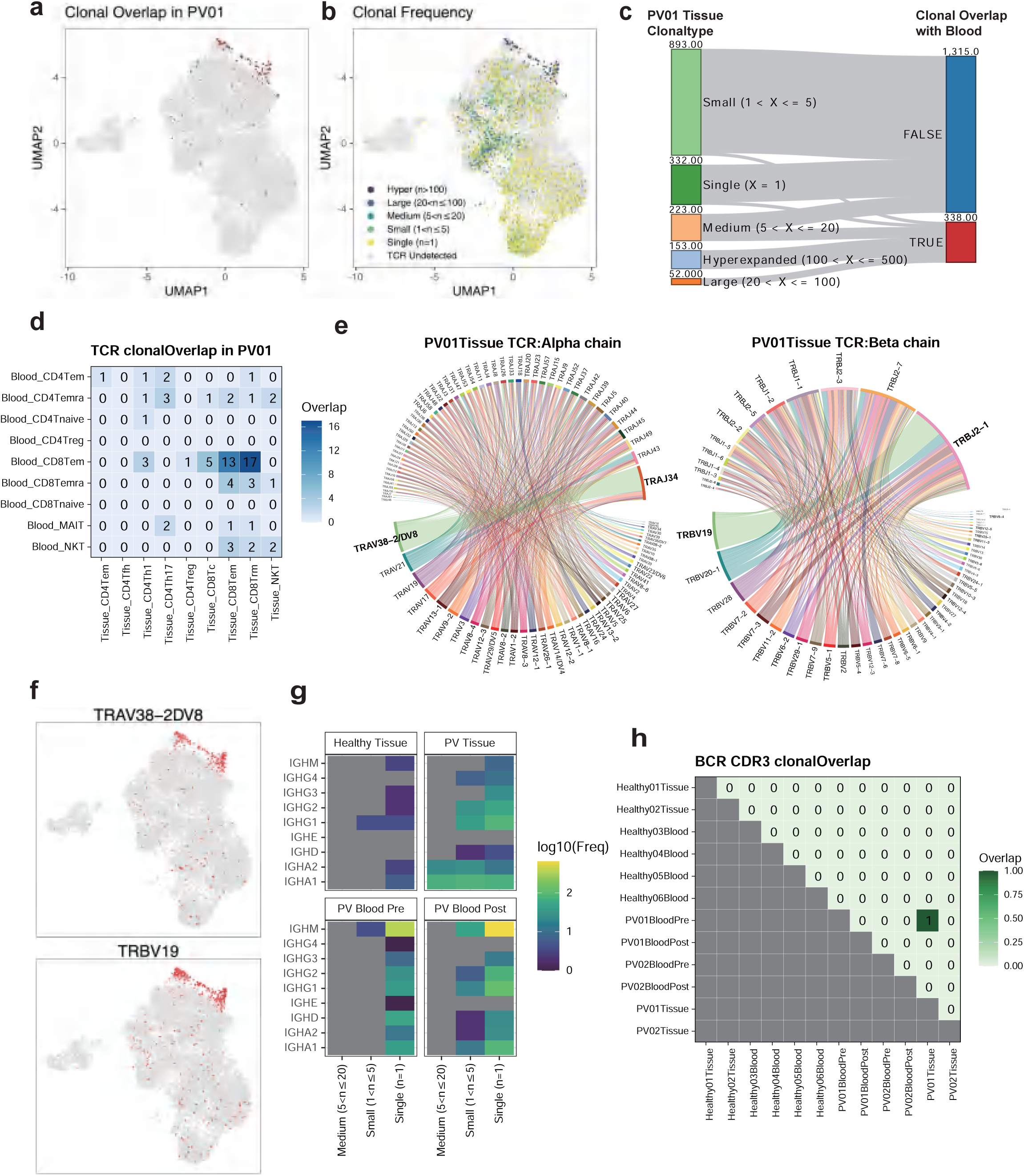
The characteristics of TCR and BCR in PV. **a** T cells in PV01 tissue which had clonotype overlap with peripheral blood before treatment. **b** UMAP of T cells overlaid with levels of clonal frequency. **c** Sankey plot shows whether T cells in PV01 tissue with different clonal frequency had clonal type overlap with T cells in peripheral blood. **d** Sharing of TCR clonotypes between PV01 patient’s blood before treatment and tissue. **e** Chord diagrams exhibit usage of V and J gene pairs in TCR’s alpha chains and beta chains in different samples. Links between genes indicate the frequencies of the gene pairs. **f** Feature plots show “TRAV38−2DV8” and “TRBV19” genes were both expressed in a certain group of cells. **g** Heatmap displays the clonal frequency of B cells with kinds of immunoglobulin types’ BCR in groups. **h** Sharing of BCR CDR3 sequences between blood and buccal mucosa tissues across all PV patients and healthy controls.

**Fig. 8.**
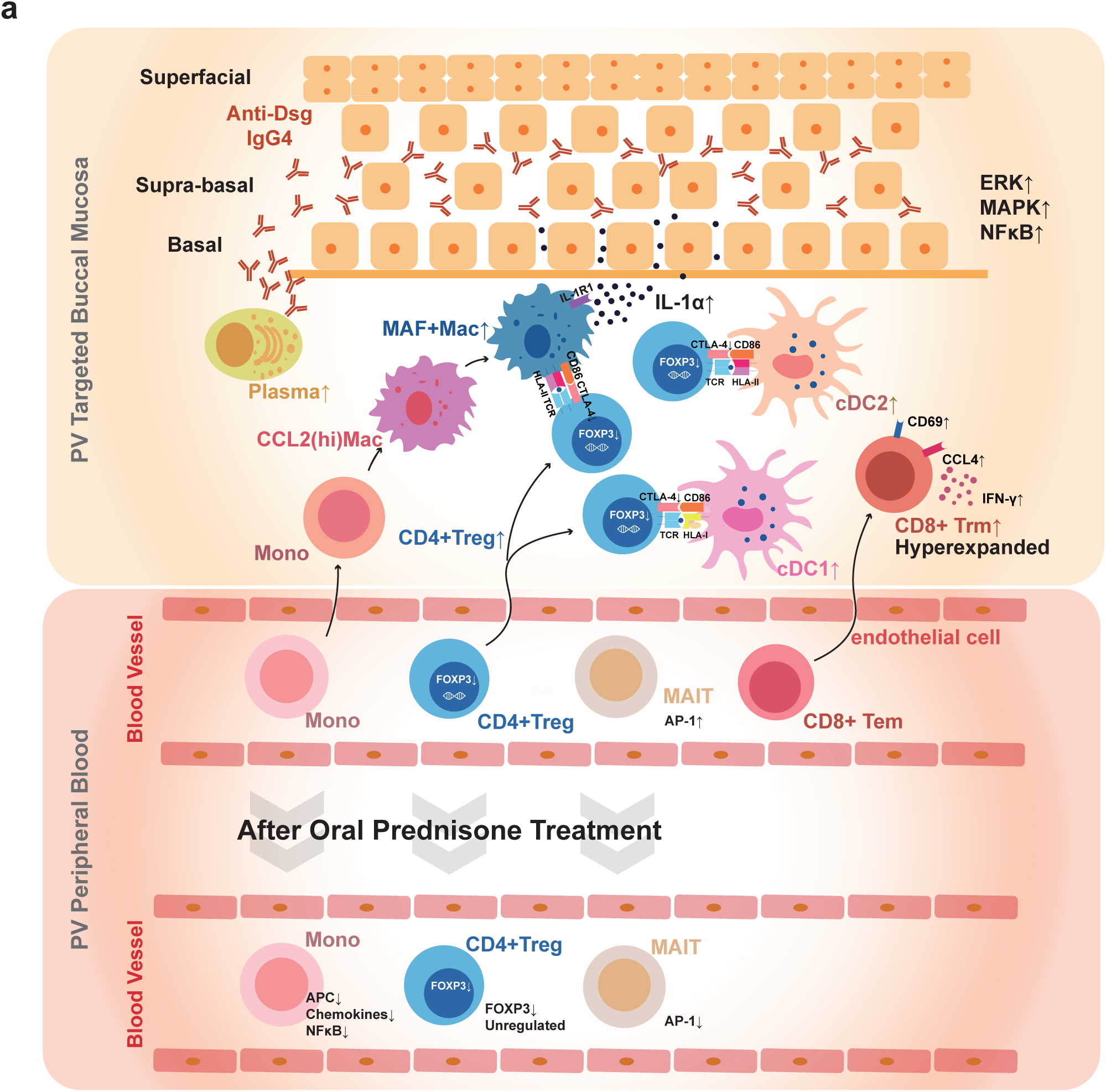
Schematic diagram summarizes the full-text.

As PV has been identified as being caused by humoral autoimmune response, we also analyzed the BCR sequence combined with scRNA-seq data. It has been pointed out that IgG4 were produced during active phase of the disease, and they were absent in patients in remission[77]. Thus, IgG4 are considered to display most of pathogenic properties [78]. Our results are consistent (Fig. 7g). There is a BCR clonotype overlap in PV01 patient’s lesion and peripheral blood (Fig. 7h). This clonotype may be the autoantibody clonotype produced by PV01 patient against the disease autoantigen. Its complementarity-determining regions (CDR)3 amino acid sequence is CAKYLYDHSFYGLDVW_CLQTLQPPYTF.

## DISCUSSION

Our study systematically compared the differences in cell type composition, gene expression, and cell-cell interactions between PV patients and HCs. Furthermore, we compared the gene expression and transcription factor alteration profiles in peripheral blood of PV patients before and after oral prednisone therapy that cured all lesions. Our findings are consistent with previous studies, and we also discovered some interesting and surprising results.

Based on our results, we observed that various types of immune cells were activated in the peripheral blood and enriched in the targeted tissue to engage in the local immune response of PV. We also observed that multiple signaling pathways, including cytokine-mediated, ERK, MAPK, and NFκB, were increased in the adjacent epithelium of PV erosions. Additionally, the IL-1 signaling pathway among epithelial subclusters and myeloid subclusters was up-regulated, and IL1B expression was up-regulated in myeloid subclusters in peripheral blood of PV and down-regulated after treatment. Combining our data with previous studies [44, 45, 55], we indicate IL-1α was released from damaged epithelial cells and sensed by infiltrating myeloid cells, thus leading to the recruitment of more inflammatory myeloid cells to the site of tissue damage. And we suggest that systemic and local IL-1 inhibition has the potential to be an effective complementary or alternative therapy for PV. Furthermore, we observed that myeloid cells could be divided into several subsets based on specific gene expression patterns, and they were likely to perform different duties such as cell chemotaxis and antigen presentation. We found that these roles, especially the antigen presentation role, were enhanced in PV, whether in the blood or local lesions.

We found that the proportion of CD8+ Trm cells in PV lesions was higher compared to HCs, whereas current literature has primarily focused on the contributions of CD4+ Trm cells to the severity and refractoriness of pemphigus and their role in local immunological pathogenesis [29]. Additionally, CD8+ Trm cells in PV lesions exhibited higher expression of tissue residency and T cell activation-related genes such as CD69[79], inflammatory cytokines CCL4 and IFNG, than healthy controls. Furthermore, their interaction with other cells was significantly enhanced in IFN-II signaling pathways in PV lesions. Surprisingly, we also found high clone intensity of CD8+ Trm cells in lesions and their connection with CD8+ Tem cells in blood through combining TCR sequencing results with RNA sequencing data. Collectively, these findings suggest a potential role for CD8+ Trm cells in the pathogenesis of PV. CD4+ Treg cells were found to be dysfunctional in both blood and lesions of PV patients, as previous studies have shown[9–15]. The low expression of FOXP3 in CD4+ Treg cells in peripheral blood may lead to weakened inhibitory effects on immune cells. Furthermore, the weakened inhibitory interaction between CD4+ Treg cells and myeloid subclusters may also contribute to autoimmunity overreaction in PV. Interestingly, the expression level of FOXP3 in CD4+ Treg cells in PV’s blood could not be improved even after oral prednisone treatment successfully cured all lesions. However, various pro-inflammatory transcription factors and genes were down-regulated after treatment. Prednisone treatment’s regulatory effect on PV was cell-specific, as monocytes and MAIT cells were in a state of high inflammation before treatment and had the greatest changes after all lesions were cured. These two cell types may be highly related to the inflammation and autoimmune activity of PV, and they may also be important specific targeted cells of prednisone. Nevertheless, the inflammatory transcription factors of monocytes were still at a high level after treatment, which may be related to the chronic state of PV.

## CONCLUSIONS

In summary, our study provides new insights into the roles of immune cell subtypes and epithelial cells in the pathology of PV, based on the analysis of peripheral blood and buccal mucosa samples. We identified antigen-presenting myeloid cells, inflammatory CD8+ Trm, and dysfunctional CD4+ Treg as key players in PV. We also found up-regulated signaling pathways in epithelial cells, with IL-1 signaling being a promising therapeutic target. Our results suggest that prednisone has cell-type-specific effects, regulating MAIT and monocytes more strongly than CD4+ Treg. Moreover, we provide CDR3 amino acid sequence data of BCR that may serve as potential therapeutic targets. Although our study didn’t reveal the trigger of the autoimmune response in PV, it provides a comprehensive framework for understanding the pathology of PV, particularly the mucosal-dominant type.

## DATA AVAILABILITY

Raw sequencing reads of the scRNA-seq and scTCR/BCR-seq experiments generated for this study have been deposited in Genome Sequence Archive (GSA)-Human database (identifier HRA003753).

## ACKNOWLEDGEMENTS

This work is supported by the National Natural Science Foundation of China (No. U19A2005, 82270986, 81972551, 81730030), the CAMS Innovation Fund for Medical Sciences (CIFMS) (2020-I2M-C&T-A-023, 2019-I2M-5-004), and Natural Science Foundation of Sichuan Province (2022NSFSC0054), and the Young Elite Scientist Sponsorship Program by CAST (2021QNRC001).

## AUTHOR CONTRIBUTIONS

Xin Zeng and Taiwen Li participated in the study conceptualization and experimental design. Shumin Duan, Taiwen Li and Qionghua Li participated in data analysis. Shumin Duan drafted the manuscript. Taiwen Li revised the article. Xin Zeng and Taiwen Li helped with implementation. All authors contributed to refinement of the study protocol and approved the final manuscript.

## COMPETING INTERESTS

All authors declare no competing interests.

---

**Fig. S1 a Clinical pictures of 2 PV patients.** Clinical pictures show 2 patients had erosions affecting multiple sites of oral mucosa. The lesions were cured after oral prednisone treatment. **b** UMAP plot of epithelial cells in tissue before correction by Harmony(left panel), and after correction by Harmony(right panel). Color-coded for the sample origins. **c** UMAP plot of myeloid cells in tissue before correction by Harmony(left panel), and after correction by Harmony(right panel). Color-coded for the sample origins. **d** Proportion of myeloid subclusters in healthy tissues and PV tissues.

**Fig. S2 a** UMAP plot of T/NK cells in tissue before correction by Harmony(left panel), and after correction by Harmony(right panel). Color-coded for the sample origins. **b** UMAP plot of immune cells in blood before correction by Harmony(left panel), and after correction by Harmony(right panel). Color-coded for the sample origins. **c** Dot plot shows gene enrichment analysis of up-regulated differential expression genes of monocytes in PV blood before treatment(PV Blood Pre) in comparison with healthy controls(left panel). Dot plot shows gene enrichment analysis of down-regulated differential expression genes of monocytes in PV blood after treatment(PV Blood Post) in comparison with PV blood before treatment(PV Blood Pre) (right panel).

**Fig. S3 a** Sankey plot shows whether T cells in PV01 blood before treatment with different clonal frequency had clonal type overlap with T cells in PV01 tissue. **b** Sankey plot shows whether T cells in PV02 tissue with different clonal frequency had clonal type overlap with T cells in PV01 blood before treatment. **c** Sankey plot shows whether T cells in PV02 blood before treatment with different clonal frequency had clonal type overlap with T cells in PV02 tissue. **d** Sharing of TCR clonotypes between PV02 patient’s blood before treatment and tissue.

## Notes

### Competing Interest Statement

The authors have declared no competing interest.

