## supplementary table and figure for "Single-Cell Transcriptomes and Immune Repertoires Reveal the Cell State and Molecular Changes in Pemphigus Vulgaris"

**Table S1: Clinical features of enrolled individuals**

| Patient ID | Gender | Age (years) | Course of disease in first visit (months) | Weight （kg） | PDAI | Nikolsky sign | Anti-Dsg1 （μ/ml） | Anti-Dsg3 （μ/ml） | Dosage of prednisone （mg/day） | Course of tapering of prednisone (days) |
|---|---|---|---|---|---|---|---|---|---|---|
| PV01-pre | Female | 57 | 24 | 52 | 17,20 | + | 34.33 | 167.17 | 45 | 90 |
| PV01-post | | | | NA | 0 | NA | 9.40 | 124.37 | 30 | |
| PV02-pre | Female | 32 | 6 | 52.5 | 21,27 | + | 25.43 | 252.31 | 60 | 66 |
| PV02-post | | | | NA | 0 | NA | 20.48 | 172.67 | 40 | |
| HC01 | Male | 24 | NA | NA | NA | NA | NA | NA | NA | NA |
| HC02 | Male | 21 | NA | NA | NA | NA | NA | NA | NA | NA |

NA: not available; PV: pemphigus vulgaris; HC: healthy controls

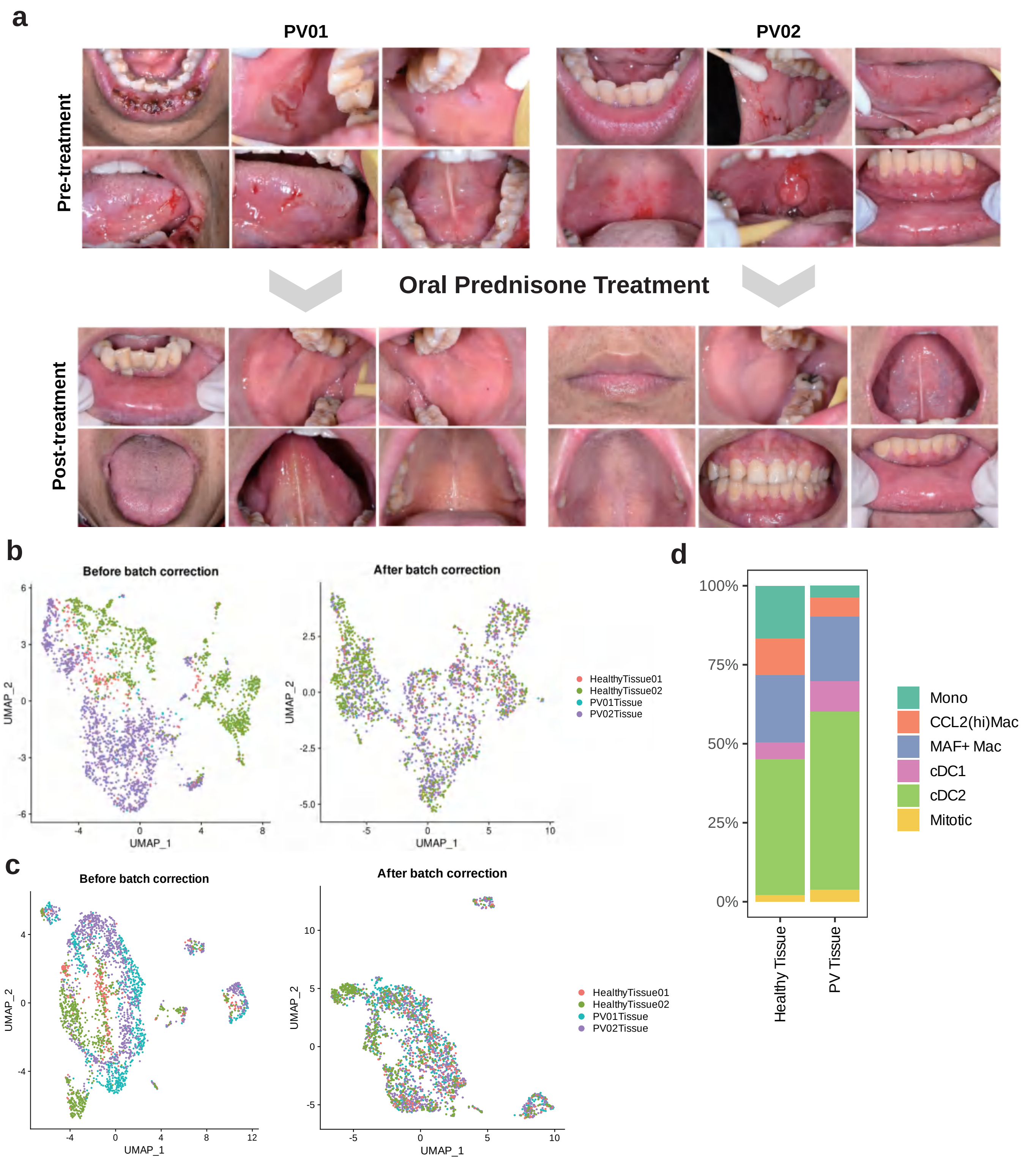
a
PV01
PV02
Pre-treatment
Oral Prednisone Treatment
Post-treatment
b
Before batch correction
After batch correction
UMAP_1
UMAP_2
HealthyTissue01
HealthyTissue02
PV01Tissue
PV02Tissue
c
Before batch correction
After batch correction
UMAP_1
UMAP_2
HealthyTissue01
HealthyTissue02
PV01Tissue
PV02Tissue
d
100%
75%
50%
25%
0%
Healthy Tissue
PV Tissue
Mono
CCL2(hi)Mac
MAF+ Mac
cDC1
cDC2
Mitotic

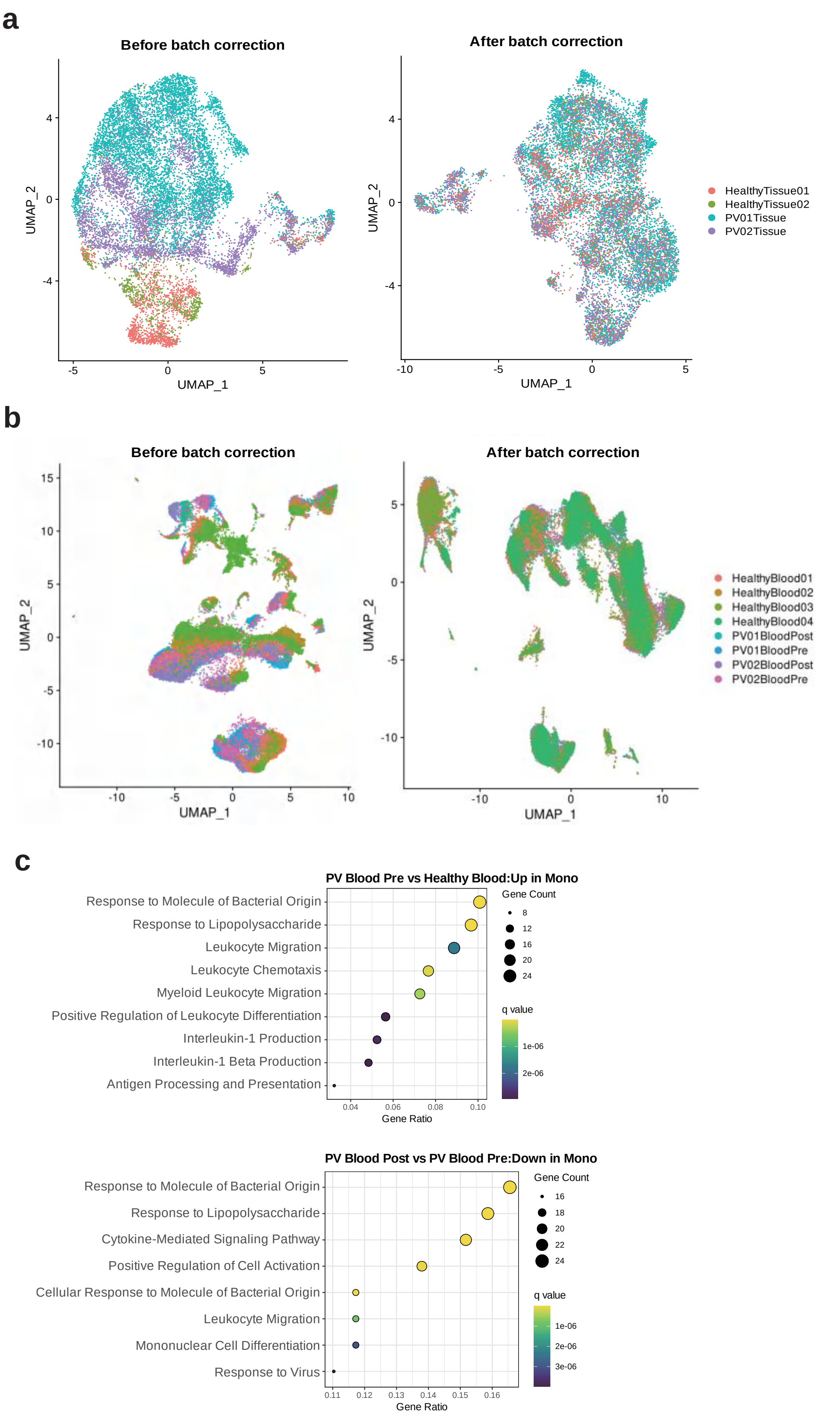

a
Before batch correction
After batch correction
UMAP_2
UMAP_1
4
0
-4
-5
0
5
-10
HealthyTissue01
HealthyTissue02
PV01Tissue
PV02Tissue
b
Before batch correction
After batch correction
15
10
5
0
-5
-10
UMAP_2
UMAP_1
HealthyBlood01
HealthyBlood02
HealthyBlood03
HealthyBlood04
PV01BloodPost
PV01BloodPre
PV02BloodPost
PV02BloodPre
c
PV Blood Pre vs Healthy Blood:Up in Mono
Response to Molecule of Bacterial Origin
Response to Lipopolysaccharide
Leukocyte Migration
Leukocyte Chemotaxis
Myeloid Leukocyte Migration
Positive Regulation of Leukocyte Differentiation
Interleukin-1 Production
Interleukin-1 Beta Production
Antigen Processing and Presentation
0.04
0.06
0.08
0.10
Gene Ratio
Gene Count
8
12
16
20
24
q value
1e-06
2e-06
PV Blood Post vs PV Blood Pre:Down in Mono
Response to Molecule of Bacterial Origin
Response to Lipopolysaccharide
Cytokine-Mediated Signaling Pathway
Positive Regulation of Cell Activation
Cellular Response to Molecule of Bacterial Origin
Leukocyte Migration
Mononuclear Cell Differentiation
Response to Virus
0.11
0.12
0.13
0.14
0.15
0.16
Gene Ratio
Gene Count
16
18
20
22
24
q value
1e-06
2e-06
3e-06

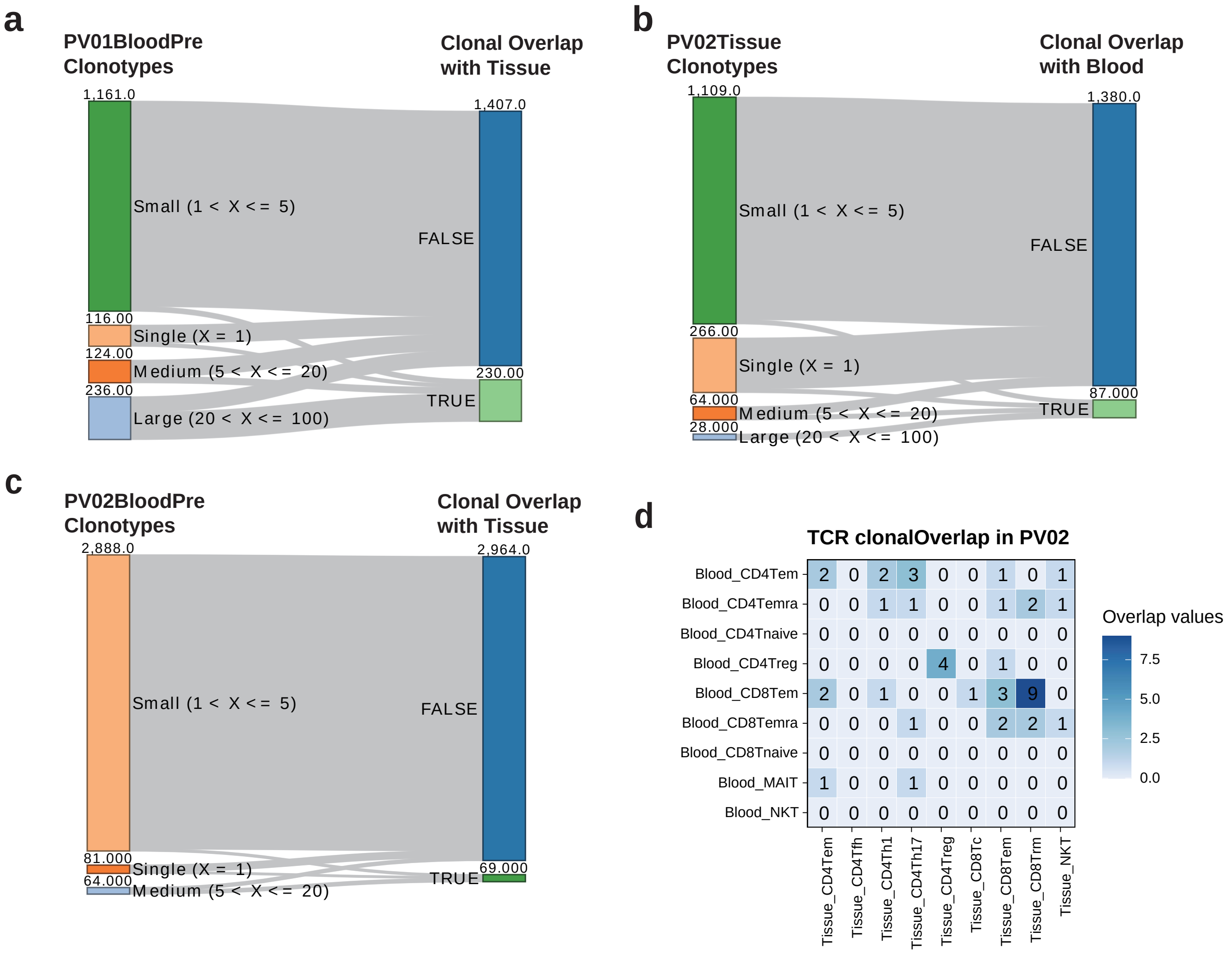
a
PV01BloodPre Clonotypes
Clonal Overlap with Tissue
1,161.0
Small (1 < X <= 5)
116.00
Single (X = 1)
124.00
Medium (5 < X <= 20)
236.00
Large (20 < X <= 100)
1,407.0
FALSE
230.00
TRUE
b
PV02Tissue Clonotypes
Clonal Overlap with Blood
1,109.0
Small (1 < X <= 5)
266.00
Single (X = 1)
64.000
Medium (5 < X <= 20)
28.000
Large (20 < X <= 100)
1,380.0
FALSE
87.000
TRUE
c
PV02BloodPre Clonotypes
Clonal Overlap with Tissue
2,888.0
Small (1 < X <= 5)
81.000
Single (X = 1)
64.000
Medium (5 < X <= 20)
2,964.0
FALSE
69.000
TRUE
d
TCR clonalOverlap in PV02
Blood_CD4Tem
Blood_CD4Temra
Blood_CD4Tnaive
Blood_CD4Treg
Blood_CD8Tem
Blood_CD8Temra
Blood_CD8Tnaive
Blood_MAIT
Blood_NKT
Tissue_CD4Tem
Tissue_CD4Tfh
Tissue_CD4Th1
Tissue_CD4Th17
Tissue_CD4Treg
Tissue_CD8Tc
Tissue_CD8Tem
Tissue_CD8Trm
Tissue_NKT
2 0 2 3 0 0 1 0 1
0 0 1 1 0 0 1 2 1
0 0 0 0 0 0 0 0 0
0 0 0 0 4 0 1 0 0
2 0 1 0 0 1 3 9 0
0 0 0 1 0 0 2 2 1
0 0 0 0 0 0 0 0 0
1 0 0 1 0 0 0 0 0
0 0 0 0 0 0 0 0 0
Overlap values
7.5
5.0
2.5
0.0
